# AI-Assisted Cryo-ET Workflow for 3D Visualization of Chromatin during Cellular Differentiation

**DOI:** 10.1101/2025.10.15.682601

**Authors:** John Watt, Cullen Roth, Isaac Huber, Alexander S. Hall, Manish Kumar Singh, Sofiya Micheva-Viteva, Karissa Y. Sanbonmatsu, Shawn R. Starkenburg, Christina R. Steadman

## Abstract

Understanding how chromatin architecture changes during cellular differentiation requires structural methods that can resolve native genomic organization at high resolution. Here, we present an AI-assisted cryo-electron tomography (cryo-ET) and segmentation workflow to quantify chromatin compaction across various stages of motor neuron differentiation from induced pluripotent stem cells (iPSC). By directly imaging extracted and vitrified chromatin, we preserve native structure and avoid artifacts from heavy metal staining and resin embedding. Using three-dimensional (3D) density analysis, we measure chromatin density and capture the progressive increase in chromatin compaction with lineage commitment. This is then correlated with population-averaged Hi-C experiments, observing consistency between the microscale higher order structure of chromatin and global contact patterns. Our approach enables direct visualization of chromatin organization under near-physiological conditions, bridging the gap between structural imaging and genome-wide contact mapping. This platform therefore establishes an AI-assisted experimental framework for linking chromatin architecture to regulatory mechanisms during differentiation.

## Introduction

In nature, genomic DNA does not exist as free and stable linear strands. Rather, the genome is spatially organized with linear DNA molecules tightly associated with histone proteins forming the nucleosome, which is the fundamental unit of chromatin (1). The nucleosome is defined as 147 base pairs of DNA nucleotides wrapped around a core histone octamer, a squat cylinder approximately 11 nm in diameter and 5.5 nm in height (2). Separated by linker DNA, nucleosomes can then interact with one another and condense into a compact fiber, which is further organized into higher order structures that allow the genome to efficiently pack into the nucleus within the boundaries of the nuclear envelope. The textbook view of chromatin packing is that 10 nm chromatin fibers assemble into 30-nm fibers, that further fold into 120-nm chromonema, to 300-to 700-nm chromatids and finally, into mitotic chromosomes (3,4). However, the condensed 30 nm chromatin fiber is potentially an artifact of sample isolation in high salt concentration conditions and not the predominant form of chromatin organization in cells (5,6). A recent study showed that during both interphase and mitosis, chromosome fibers consisted of disordered chains with diameters between 5 and 24 nm (7). Although debate continues over the precise nature of chromatin organization beyond the nucleosome, it is well established that higher-order structure plays a central role in regulating gene accessibility and transcriptional activity (8-10).

Higher-order chromatin structures have been characterized across a wide range of length scales. These include larger domains identified as open (active gene expression) A compartments and closed (repressive gene expression) B compartments, which can be further partitioned into smaller sub-domains, known as topologically associating domains (TADs). TADs are defined as regions that exhibit a high frequency of intra-chromosomal interactions—containing features such as chromatin loops—and relatively few interactions with regions outside their respective compartments (11). Such structural features define regions of transcriptional activity and play a critical role in gene expression, transcription, and repair by providing a steric barrier to DNA access (12). Further, chromatin is structurally dynamic and throughout the cell cycle, chromatin reversibly alternates between closed and accessible configurations, the balance of which ultimately determines cell fate (13). While many studies have structurally characterized chromatin both *in vivo* and *in vitro*, uncertainty remains about the nature and evolution of higher-order chromatin structure, as well as the structural and physical mechanisms driving compaction (14). To improve this understanding, we require precise methods the preserve native chromatin characteristics and accurate quantification strategies.

Early models of chromatin organization were limited by imaging resolution; however, advances in X-ray diffraction and electron microscopy (EM) have recently provided more insight. While useful structural information has been gleaned from studies employing these advanced techniques, they relied on preserving or fixing the samples using harsh chemicals, dehydration steps, and staining (15,16). Such harsh conditions influence the native sample configuration and introduce structural changes. Alternatively, cryogenic electron tomography (cryo-ET), which facilitates high-resolution, three-dimensional imaging, can be used to characterize chromatin in its native hydrated state, offering unprecedented clarity into native chromatin organization. For example, Hou et. al. used cryo-FIB and cryo-ET to investigate the 3D architecture of native chromatin in human T-lymphoblasts during intact interphase. They found that most nucleosomes are connected by straight linker DNA, forming a flexible and relaxed zig-zag pattern, rather than the expected condensed 30-nm fiber (4). Hatawaza et. al. developed a crosslinking method to isolate native nucleosomes from human cells, enabling structural determination of nucleosomes and chromatin subunits using cryo-EM and cryo-ET (6).

Whereas cryo-EM directly visualizes chromatin density and fiber organization at nanometer resolution, chromosome conformation capture techniques such as Hi-C, have emerged as indispensable tools for mapping genome-wide spatial interactions between distal DNA elements, and elucidating their effect on gene regulation. Hi-C works by measuring the frequency (as an average over a cell population) at which two DNA fragments physically associate in 3D space, linking chromosomal structure directly to the genomic sequence. The general procedure of Hi-C involves first crosslinking chromatin material using formaldehyde. Then, the chromatin is solubilized and fragmented, and interacting loci are re-ligated together to create a genomic library of chimeric DNA molecules. The library is then sequenced and produces reads indicating the relative abundance of these chimeras, or ligation products; this is correlated to the probability that the respective chromatin fragments interact in 3D space across the cell population (17). While Hi-C offers powerful insights into chromatin interactions, it does not produce a direct visualization of higher-order structure.

By integrating Hi-C’s genome-wide contact maps derived from sequencing with cryo-ET’s direct visualization of chromatin fibers, population-averaged interaction frequencies can be connected to the underlying physical structure of higher-order chromatin. Cai et al. used cryo-electron tomography (cryo-ET) to visualize the three-dimensional arrangement of nucleosomes and other macromolecular complexes within *Schizosaccharomyces pombe* (yeast) during interphase and mitosis. They reported nucleosome clusters visualized by cryo-ET are smaller than expected for Hi-C domains, highlighting a scale mismatch, based on comparisons with previous Hi-C results found in the literature (18). By pairing chromatin scanning transmission electron tomography (ChromSTEM) with multi-omic analysis and single-molecule localization microscopy, Li et. al. studied the role of cohesin in regulating the conformationally defined chromatin nanoscopic packing domains. Their results showed that the 3D genomic structure, composed of packing domains, is generated through cohesin activity and nucleosome modifications (19). However, these samples were prepared from cancer cells with a buffer containing a multivalent cation, which is known to induce chromatin compaction, and involved multiple dehydration, staining and embedding steps, meaning the chromatin was analyzed far from its native hydrated state. While these seminal results have broadened our foundational understanding of chromatin structure, the methods employed, particularly for preparation of chromatin molecules for visualization, may confound ground truth estimates of definitive chromatin structure.

During cellular differentiation, chromatin undergoes dynamic structural rearrangements in which genomic regions that do not contribute to the cell lineage specification compact into transcriptionally silent heterochromatin, while genes defining the cell identity remain accessible as euchromatin. These changes establish and maintain the cell-type–specific gene expression patterns that underlie specialized function. Dixon et. al. mapped genome-wide chromatin interactions in human embryonic stem (ES) cells and four differentiated lineages derived from ES cells, revealing widespread chromatin reorganization associated with lineage specification (20). Given these profound chromatin changes that occur during maturation, we investigate the evolution of chromatin compaction during the differentiation of inducible pluripotent stem cells (iPSCs). We employ microstructure analysis from cryo-ET with methods to maintain native chromatin configurations and global contact frequency analysis from Hi-C experiments. We develop an AI-assisted framework to correlate cryo-ET visualizations with Hi-C contact maps to determine the impact of native chromatin compaction that occurs during differentiation of iPSCs into human neuronal stem cells (NSCs) and ultimately human motor neurons (MNs). By applying correlated cryo-ET and Hi-C analyses across this differentiation pathway, we quantify chromatin conformation shifts from its highly dynamic and accessible state to a more compartmentalized and topologically constrained structure.

## Materials and Methods

### Cell culture and differentiation

Human iPSCs (WTC-11 cell line, Coriell Institute) were cultured and maintained in Gibco Essential 8™ medium (Thermo Fisher Scientific). iPSC differentiation into motor neurons was previously described (21).

### Hi-C protocol for *in situ* chromatin conformation capture

The Arima Hi-C Kit User Guide Mammalian Cell Lines Protocol (Arima Genomics, A5100008) was applied for Hi-C sample preparation. Cells (10^6^) were cross-linked with methanol-free formaldehyde (final concentration 2%) for 10 min at RT with continuous mixing. Stop solution (from Arima kit A5100008) was added for 5 min at RT followed by 10 min incubation on ice. Cells were pelleted by centrifugation, washed with PBS, and distributed in 10^6^ cell aliquots. Cell pellets were stored at -80°C until further processing. DNA was purified with AMPure XP beads and sheared to 400 bp fragments using the Covaris Focused-Ultrasonicator instrument. On-bead libraries were prepared for Illumina NextSeq2000 with the NEBNext Ultra II DNA Library Prep kit (NEB, E7645L) following the manufacturer’s protocol.

### Chromatin extraction for cryo-ET

Chromatin was extracted from cells using a modified protocol (22,23). First, salicylic acid coated magnetic nanoparticles were synthesized by adding a 2:1:4 molar ratio of FeCl3·6H2O:FeCl2·4H2O:salicylic acid to a three-neck flask containing NaOH solution (pH 11.0) with vigorous stirring with N2 gas. After refluxing at 90 ^o^C for 4 h, a dark brown suspension was formed. The magnetic nanoparticles were then separated with a magnet and washed with DI water, before being dispersed into to 10 mg/mL suspension in DI water for use. Prior to chromatin extraction, cells (10^6^) were fixed with 2% methanol-free paraformaldehyde for 10 min followed by rapid neutralization and removal of the fixing agent via centrifugation. Cell pellets were collected in an Eppendorf tube by centrifugation and lysed with 100 *μ*L of lysis buffer (250 mM SDS, 1 mM EDTA, 0.5 mM EGTA, 1 mM PMSF and 2% Protease Inhibitor Cocktail). To disperse, the cell pellet was pipetted up and down slowly, then 10 *μ*L of 10 mg/mL magnetic nanoparticle suspension was added, and the solution was incubated for 10 min. Isopropanol was added to form a complex between chromatin and the magnetic nanoparticles, and the solution was incubated for another 5 min. The chromatin-NPs complexes were then isolated by a magnet and washed once with PBS buffer. Chromatin was eluted away from the magnetic NPs in 50 mL of PBS buffer and incubated for 4 h at RT or overnight at 4 ^o^C. Previous studies included 1 mM MgCl2 in this elution step; however, this has been shown to induce artificial chromatin condensation and was omitted here (24). The magnetic nanoparticles were then separated by a magnet, leaving the released chromatin suspended in the supernatant. The concentration of chromatin was determined from DNA absorbance at 260 nm by an Eppendorf Biospectrometer.

### Cryogenic Electron Tomography (cryo-ET)

Vitrification for cryo-ET was then performed using a Thermo Fisher Scientific Vitrobot Mark IV™ with a manual back blotting step in place of the automatic blotting procedure. Here, the automated settings (Blot Time, Drain Time, etc.) were all set to zero. Protochips cryo-TEM grids were made hydrophilic by plasma charging for 30 seconds. The grid was placed in the tweezer assembly and retracted into the chamber at 100% humidity. The process was started and 4 µL of sample solution pipetted onto the grid and allowed to settle for 15 sec. The door was opened, and the grid back blotted twice manually using filter paper. The first back blot removed a visible amount of liquid, whereas the second was less visible but essential to form an electron transparent vitreous ice layer. Cryo-ET was performed on a Talos L120C operating at 120 kV with a Gatan 626 Holder, using Thermo Fisher Scientific Tomography 5 software, and the image stack reconstructed in Thermo Scientific Inspect 3D Software.

### Segmentation and compaction quantification

We visualized and segmented the reconstructed tomogram to quantify chromatin compaction using a modified method consisting of two interdependent components (15). First, AI-assisted segmentation tools in Thermo Scientific Avizo™ 3D Pro 2022.2 and 2023.1 software were employed to qualitatively visualize chromatin and construct a 3D explorable tomogram. The second component quantified the chromatin volume concentration of the respective tomogram and utilized ImageJ, Avizo, and Python.

Chromatin volume concentration was calculated identifying a section of immediately adjacent chromatin fibers, designated as a domain. From the central point of each domain, pixel intensities in the image representing the volume of chromatin were summed within a specified radius. The summed intensities were divided by the total analyzed volume to yield the radial chromatin volume concentration, representing the density of chromatin within the specified radius. By incrementally increasing the radius and recalculating the chromatin volume concentration, spatial changes in chromatin compaction were determined.

Pixel values from all tomogram images were averaged along the Z-axis to produce a single representative image. A Gaussian blur (radius = 5.00 pixels) was applied, followed by the Contrast Limited Adaptive Histogram Equalization (CLAHE) algorithm to denoise and improve the signal-to-noise ratio, while preserving structural detail. The processed image was masked to isolate regions containing chromatin, and an H-minima transform was applied to generate an image overlaid with chromatin domains. Chromatin domains located within 220 nm of image boundaries were excluded. The remaining domains were labeled, with each represented by a single pixel at its central coordinate.

Radial chromatin volume concentration was calculated by linking the labeled domain center image to the original tomogram. Beginning at the XY-coordinate of each domain center, pixel intensities were summed radially outward in one-pixel increments in the XY-plane across the full Z-axis of the tomogram. The radius was expanded until reaching 220 nm from the domain center. This process was repeated for all domain centers in the image. The resulting chromatin volume concentration values were plotted as a function of radial distance from each domain center. Finally, the values for all domains were averaged at each radius to produce the overall chromatin volume concentration profile for the tomogram.

### Hi-C data processing and visualization

Hi-C libraries were generated and sequenced as described above from cultures of iPSCs, NSCs, and MNs for direct comparison to cryo-ET studies. Paired-end sequencing reads from all samples were passed through our multi-omic processing pipeline, SLUR(M)-py (25). Default settings were used for processing, and sequenced reads were mapped to the human, telomere-to-telomere, reference genome (26). Post-processing, the total number of sequenced reads were randomly subsampled (without replacement) to normalize Hi-C libraries and reduce bias in interpretation due to differences in sequencing depth (across samples). A total of 23,567,000 intra-chromosomal contacts were used as the baseline level of Hi-C contacts across all three samples. Python scripting, relying on the FAN-C library, was used to conduct analysis and generate visualizations of Hi-C data at a resolution of 750 kb (27). The paired-end sequencing data (fastq files) of the three Hi-C samples are hosted on NCBI’s sequence read archive, under the BioProject PRJNA1337150 with accession numbers SAMN52179762, SAMN52179763, and SAMN52179764.

## Results and Discussion

We substantially modified previously reported protocols for chromatin extraction to preserve all long-distance chromatin interactions through crosslinking [8, 9]. Chromatin fibers become highly compact in buffers containing significant monovalent ion concentrations [10]. Therefore, we eliminated MgCl2 in our procedure to avoid artificial stimulation of chromatin condensation. To visualize chromatin condensates using cryo-ET, the sample must be sufficiently thin to allow for transmission of the electron beam. In standard cryo-EM techniques, the sample solutions are pipetted onto grids, and excess liquid is removed with filter paper to create a thin layer, which is subsequently frozen for tilt-series collection, enabling reconstruction of 3D tomograms. However, we observed that conventional blotting removed a significant amount of material, indicating chromatin preferentially resided at the air-water interface and was likely removed the filter paper wicking. Assembly at the air-water interface has previously been observed and is a known problem in single particle analysis (28,29). We therefore employed a back blotting method that avoided contact of the air-water interface with the filter paper and promoted chromatin distribution within the grid pores.

Using this approach, we extracted chromatin from inducible pluripotent stem cells (iPSC), neuronal stem cells (NSC), and motor neuron cells (MN) and performed cryo-ET analysis (Figure 1). The resulting tomograms from this analysis (Figure 2) show significant compaction of chromatin as differentiation occurs. To create these images, we employed an AI-assisted workflow for segmentation and visualization. First, the tilt series were aligned and filtered using Inspect 3D before being imported into Avizo. AI was employed to segment the chromatin against the carbon and vitreous ice background; chromatin was visualized as voxelized renderings (Figure 2a-c). These steps follow established protocols for tomographic reconstruction and visualization; however, to identify and quantify chromatin compaction and compare across different stages of cellular differentiation, the tilt series were then subjected to further analysis. The tilt series TEM data were first consolidated into a single representative image by averaging pixel values across the Z-axis, effectively collapsing the volumetric information into a two-dimensional projection (Figure 2d-f). To suppress local intensity fluctuations, a Gaussian blur with a radius of 5.00 pixels was applied, providing a smoother baseline for subsequent processing. Noise reduction and contrast enhancement were then achieved through the application of the Contrast Limited Adaptive Histogram Equalization (CLAHE) algorithm, which improved the signal-to-noise ratio while maintaining fine structural features essential for interpretation. We then identified regions of high contrast in the micrograph using FIJI ImageJ. These domains correspond to regions of enhanced electron scattering and indicate central domains of condensed chromatin, as shown in Figure S1. We quantify compaction by analyzing the density of chromatin located radially away from each domain, and averaged them across all domains, with the normalized results shown in Figure 2g-i. Here, chromatin isolated from iPSC cells showed shows an increase in volume and variability with increasing distance from the domains. An increase in volume at larger radius indicates the mass of chromatin is less consolidated at the labeled domains, indicating chromatin was more disperse and open, which is typical of pluripotency (20). Chromatin derived from NSCs (Figure 3h) showed a moderate spread of volumes at high radius, which indicates intermediate density, with volume steadily decreasing with increasing radius. We then calculated the mean gradient and normalized it against volume to quantify how chromatin compaction changes with increasing distance from the domains. This indicated that radial compaction changes by iPSC << NSC < MN. NSC and MN showed negative normalized gradients of -2.83 × 10^-4^ and -3.28 × 10^-4^, respectively, signifying that their chromatin volumes decrease with radius which is consistent with greater compaction. Conversely, iPSC shows a small positive normalized gradient of 2.65 × 10^-4^, indicating a marginal reduction in compaction away from the domains.

**FIGURE 1.**
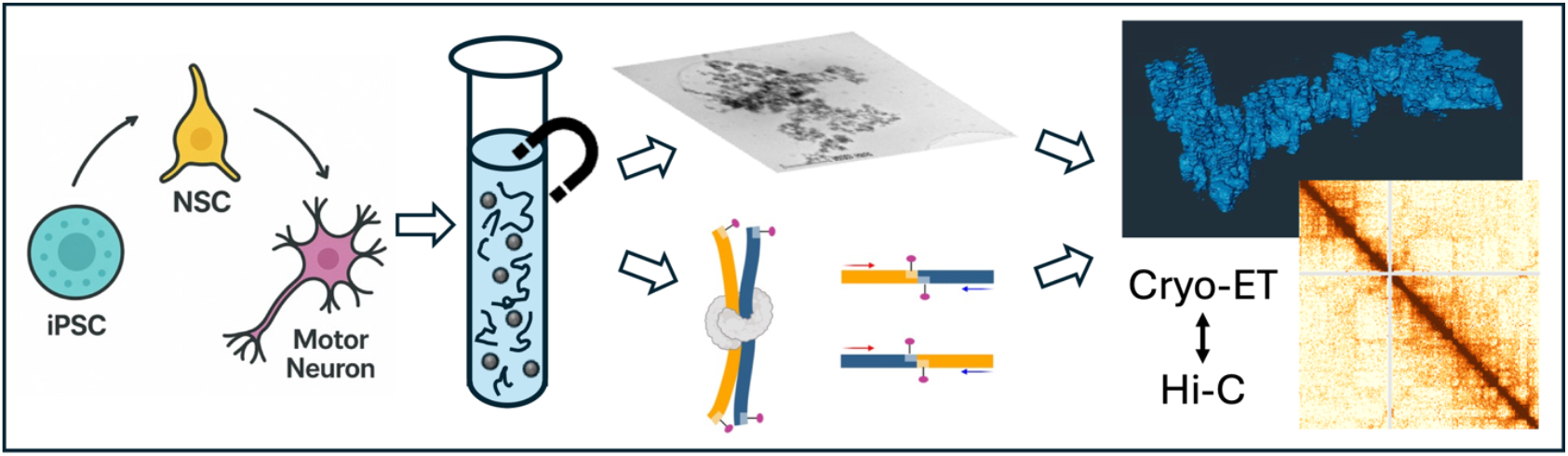
Experimental workflow to quantify chromatin compaction of cells at different stages of differentiation of inducible pluripotent stem cells (iPSC) into neuronal stem cells (NSCs) and motor neurons (MNs). Chromatin was extracted from each cell type and analyzed using cryogenic electron tomography (cryo-ET) and Hi-C to provide correlative information on the higher-order structure of chromatin.

**FIGURE 2.**
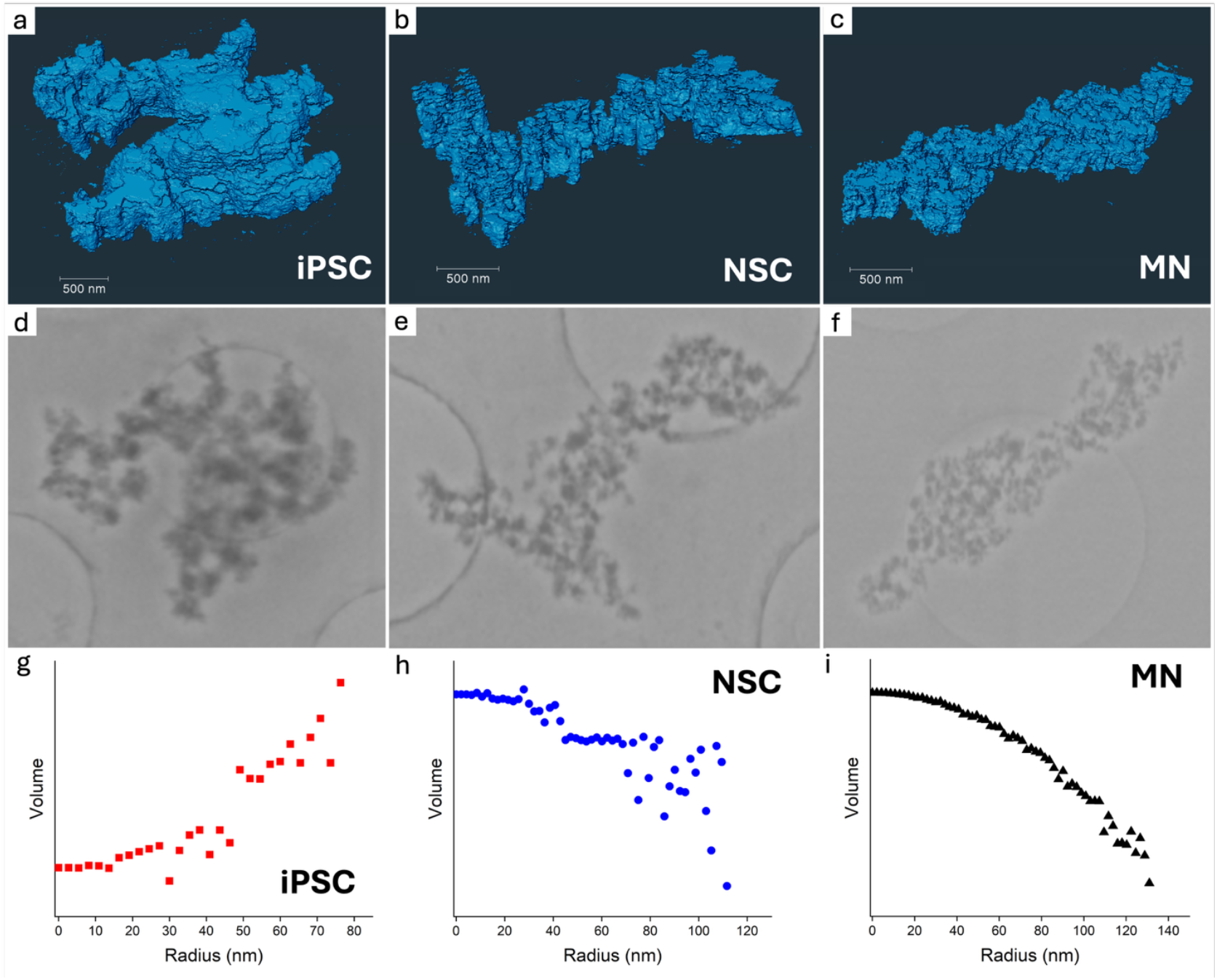
Cryo-EM and cryo-ET images from chromatin extracted from iPSC, NSC, and MN cells. **(a-c)** AI-assisted segmented and reconstructed 3D tomograms viewed in Avizo Software. (d-f) Cryo-EM images of extracted chromatin that have been filtered with a Gaussian blur and modified with contrast local adaptive histogram equalization (CLAHE) in ImageJ, to more effectively find the high contrast domain locations. (g-i) Radial volume distribution of chromatin averaged across all measured domains for iPSC, NSC, and MN cells, respectively.

**FIGURE 3.**
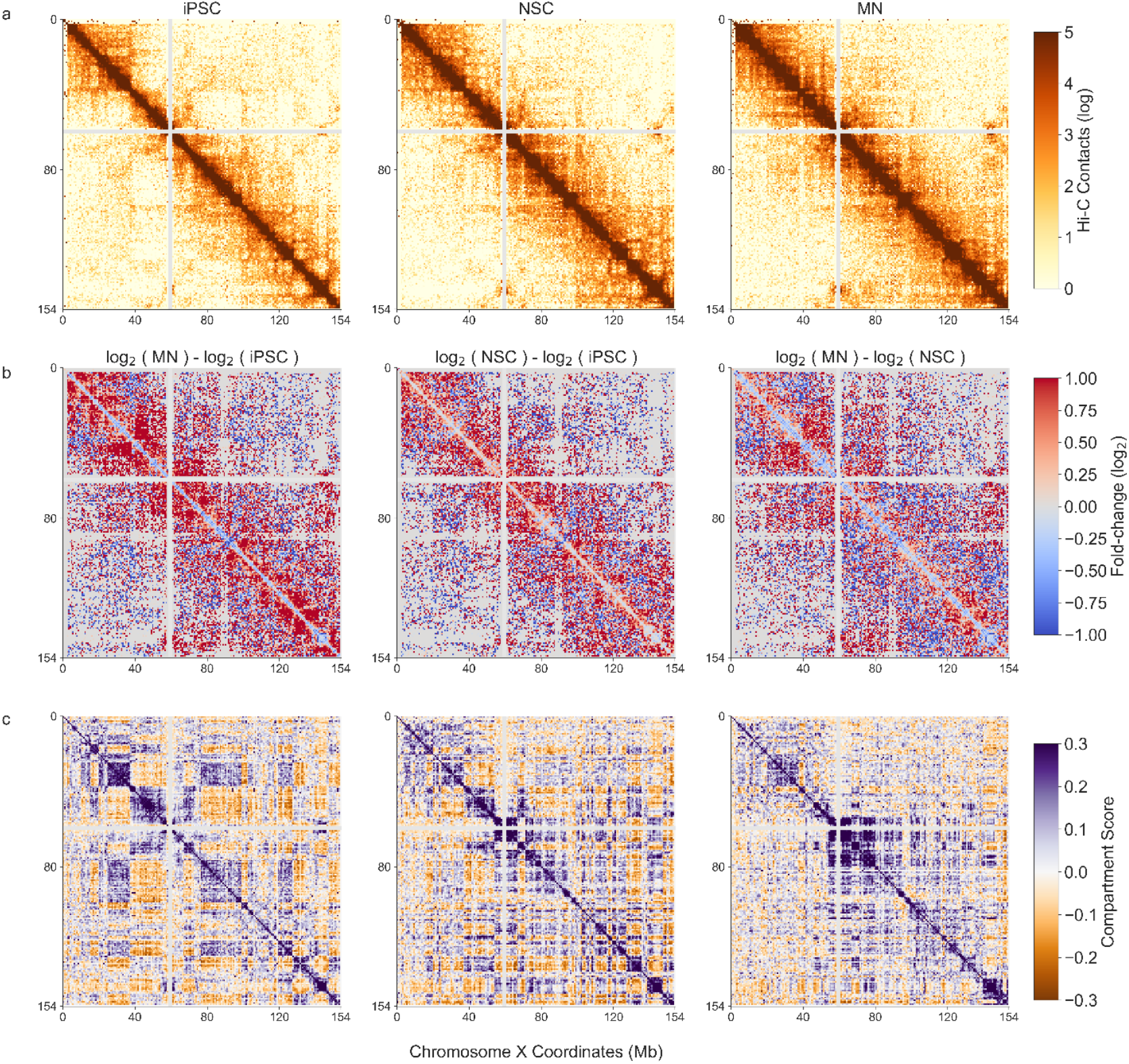
Chromatin conformation analysis across the differentiation spectra of induced pluripotent stem cells (iPSC). a) Chromosome contacts (log) at 750 kb resolution across chromosome X for iPSC, neural stem cells (NSC), and motor neuron (MN) cells (columns, left to right, respectively). Darkening orange colors indicate a higher frequency of chromatin contacts between distal chromatin regions. The diagonal space represents the linear chromosome sequence. b) The fold-change (log2) in Hi-C contacts between MN and iPSC cells, NSC and iPSC cells, and MN and NSC cells (column space, left to right, respectively). Title along each subplot annotates the direction of the fold-change relative to cell type, with red colors indicating an increasing frequency in chromosome contacts. c) Chromatin compartment scores across cell types, as in (a). Darker purple colors indicate increasing correlation in chromatin connectivity across the chromosome.

We then performed Hi-C analysis to assess if the increasing density observed in cryo-ET experiments correlated with increasing chromatin connectivity. After processing via SLUR(M)-py pipeline (25), all Hi-C data from iPSC, NSC, and MN cells was normalized to ∼23.5 million intra-chromosomal contacts to reduce visual bias between maps that emerges from differences in sequencing depth. The down-sampled Hi-C maps were used in comparisons of fold-change in contacts, chromatin compartment analysis, and visualization.

Across the stages of differentiation, we observed alterations in 3D chromatin architecture in all chromosomes (for example chromosome X, Figure 3a). The fold-change (log2) in Hi-C contacts (at 750 kb) showed increasing connectivity between linearly distal (relatively 10^6^ and 10^7^ bp apart) chromatin regions in NSC and MN cell types, when compared to iPSCs (Figure 3b). Increasing chromatin connectivity was also observed in chromatin compartments across the course of cellular development. Large alterations in chromatin compartments were observed in the transition of iPSC cells to NSC and MN (Figure 3c), with larger compartments breaking up into smaller, more tightly correlated, sub-compartments, and increasing compartmentalization across chromosome X. These patterns held across the chromosomes, with the largest changes seen in MN cells compared to iPSC or NSC cells. Taken together, these patterns indicate increasing chromatin interactions and compaction toward the linear diagonal space of the chromosomes across the stages of cellular development.

## Conclusions

The structural nature of higher-order chromatin is still under deep investigation. Here, we explored the process of chromatin compaction using the differentiation of induced pluripotent stem cells into lineage specific cell types, namely neural stem cells and motor neurons, as a model system. We employed novel methodological techniques to protect that native chromatin state during visualization with cryo-ET. We correlated density patterns calculated using AI-assisted segmentation with chromatin contact maps derived from Hi-C experiments to demonstrate a quantitative framework for investigating chromatin states. In pluripotent stem cells, the chromatin architecture is dominated by a decondensed configuration with extensive euchromatic regions, permissive to transcriptional plasticity. As differentiation progresses and stemness abates, the genome reorganizes to selectively close chromatin regions associated with pluripotency and progress toward a defined cell linage. In MNs, chromatin adopts a more compact and stable structure, with long-range interactions forming and non-neuronal regions becoming more heterochromatic and spatially segregated. NSCs represent an intermediate stage of chromatin compaction: they are distinctly more compact than the observed structure of iPSCs but remain open for determination compared to fully differentiated MNs. Together, cryo-ET and Hi-C provide a complementary view of chromatin organization, with cryo-ET resolving local microstructural features and Hi-C capturing distal genomic contacts. This integrated approach revealed how progressive compaction and spatial reorganization accompany the transition from pluripotency to a fully differentiated neuronal state.

## Acknowledgements

This material is based upon work supported by the U.S. Department of Energy, Office of Science, through the Biological and Environmental Research (BER) and the Advanced Scientific Computing Research (ASCR) programs under contract number 89233218CNA000001 to Los Alamos National Laboratory (Triad National Security, LLC) (CRS & SS). This work was further supported by the Laboratory Directed Research and Development program of Los Alamos National Laboratory under project numbers 20210134ER (CRS and KYS) and 20210082DR (KYS and SS). This work was performed, in part, at the Center for Integrated Nanotechnologies, an Office of Science User Facility operated for the U.S. Department of Energy (DOE) Office of Science.

## Author Contributions

All authors contributed to the conceptualization and design of the study. J.W., C. R., M.K.S., S.M-V. conducted the experiments and collected the data. C.R., I. H., A.S.H. analyzed the data and interpreted the results. J.W. and C.R. wrote the manuscript. K.Y.S, S.R.S, C.R.S. supervised the project and acquired funding. All authors edited, revised and approved the final version of the manuscript.

## Declaration of Interests

The authors declare no competing interests.

## Supporting Material

Supplemental information can be found online at

## Declaration of generative AI and AI-assisted technologies in the manuscript preparation process

During the preparation of this work the lead author used GhatGPT 5 to create the cartoon of cellular differentiation on the left-hand side of Figure 1. After using the tool, the authors reviewed and edited the output and take full responsibility for the content.

## Notes

### Competing Interest Statement

The authors have declared no competing interest.

