## Supporting Information for "AI-Assisted Cryo-ET Workflow for 3D Visualization of Chromatin during Cellular Differentiation"

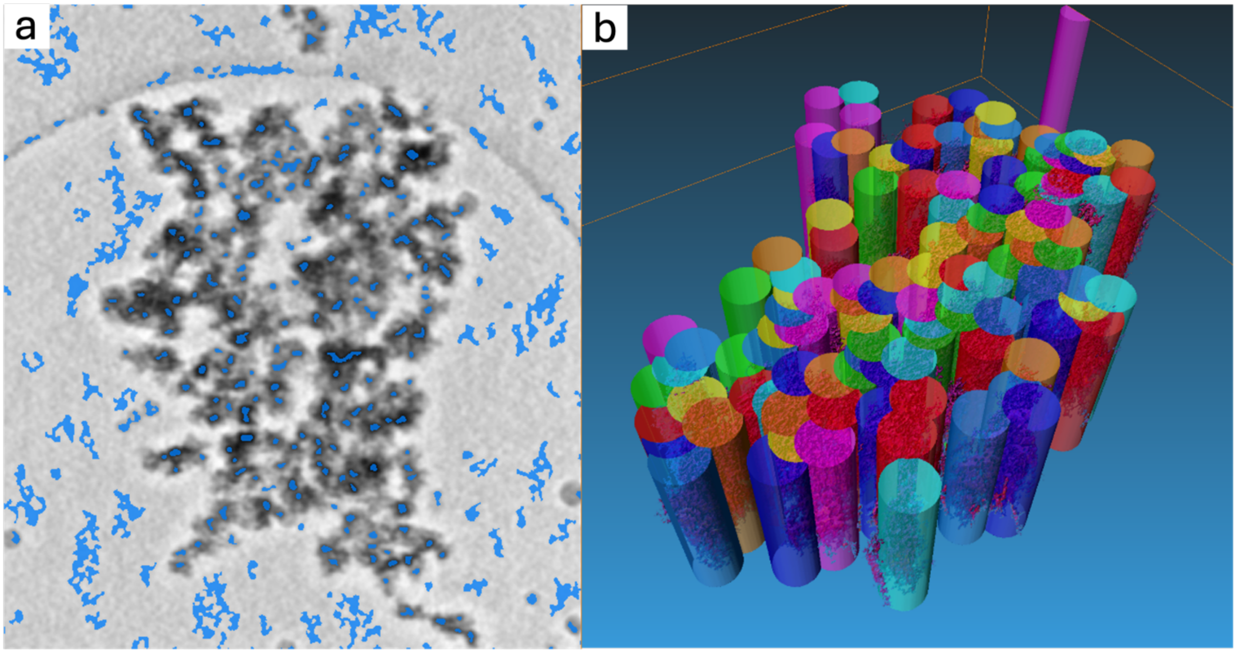


Figure S1. Identification of chromatin ‘domains’ using Image J and Thermo Scientific Avizo™ 3D Pro 2022.2 and 2023.1 software.
